# Factors Associated with Risk Of HIV-Infection Among Pregnant Women in Cameroon: Evidence from the 2016 National Sentinel Surveillance Survey of HIV and syphilis

**DOI:** 10.1101/482901

**Authors:** Jean de Dieu Anoubissi, Ekali Loni Gabriel, Cyprien Kengne Nde, Joseph Fokam, Dorine Godelive Tseuko, Arlette Messeh, Yasmine Moussa, Celine Nguefeu Nkenfou, Leonard Bonono, Serge-Clotaire Billong, Jean-Bosco Elat Nfetam

**Author notes:** Corresponding authors (JDA).

## Abstract

**Background:** Human Immunodeficiency Virus infection (HIV) remains a public health concern in Cameroon that requires regular surveillance for informed policy-making to guide programmatic interventions. Using data from the 2016 HIV national sentinel survey in Cameroon, we ascertained HIV prevalence and factors associated with risk of infection among pregnant women

**Methods:** A cross-sectional study was conducted throughout 2016 in the 10 regions of Cameroon, targeting 7000 first antenatal care (ANC-1) attendees (4000 from urban and 3000 from rural areas) in 60 sentinel health facilities. HIV serological test was performed using the national serial algorithm at the National Reference Laboratory (NRL). Prevalence was determined, and multivariate logistic regression was used to assess determinants of HIV infection, with p-value<0.05 considered statistically significant.

**Results:** Of the 7000 targeted participants, a total of 6859 first ANC-1 attendees were enrolled (98.0% sampling coverage). Median age was 26 [IQR: 21-30] years and 47,40% had a secondary school level of education. The national prevalence of HIV was 5.70% (95% CI: 4.93 – 6.40) and range from 9.7% in East region to 2.6% in North region. The prevalence was 5.58% (95% CI: 95%: 4.88 – 6.35) in urban and 5.87% (95% CI: 5.04 – 6.78) in rural settings. Factors that were associated with HIV infection included marital status, women who were married or living with their partner are less likely to be infected than singles women (aOR=0.60; 95% CI: 0.46 – 0.78), multiparity [aOR=1.5(95%CI:1.0-2.2)] and been living in the Centre, East, North-west and South-west regions. HIV infection was also significantly associated with age, with the risk of being infected increasing with age.

**Conclusion:** Pregnant women in Cameroon are still disproportionately infected with HIV compared with the general population (prevalence 4.3%). Preventive actions to curb the epidemic amongst pregnant women should prioritize interventions targeting single pregnant women, who are older, and residing particularly in the Centre, East, North West and South West regions of the country.

## Background

According to the Joint United Nations program on HIV/AIDS (UNAIDS), 36.9 million people were living with HIV in 2017 and 1.8 million were newly infected across the globe. Sub-Saharan Africa bears the greatest burden of the epidemic with 25.7 million people infected in this region[1].

Cameroon is a lower middle-income country with a generalized HIV epidemic. The country has the second largest epidemic after Nigeria in the West and Central African sub-region. The Cameroon Population-based HIV Impact Assessment (CAMPHIA) conducted in 2017 reported an HIV prevalence of 3.7% in the population aged 15-64 years[2]. This prevalence has significant disparities according to specific groups of the population and geographical areas of the country. For example, females have an HIV prevalence which is twice as high as that of males (5% and 2.3% respectively) [2] while women who are pregnant, are even more disproportionately infected by the virus.

In 2012, the national sentinel surveillance survey of HIV and syphilis (SSS) among pregnant women attending first antenatal clinic consultation (ANC1) found that 7.2% of the attendees were infected with HIV [3]. Since mother-to-child transmission (MTCT) of HIV is the leading cause of infection among children, this high prevalence among pregnant women translates to a higher risk of infection to children. Indeed, the HIV prevalence amongst HIV-exposed infants was 8.4% nationwide in that same year. In order to address this high MTCT rate of HIV, the country developed a national plan to eliminate MTCT(eMTCT plan) of the virus, through four priority approaches, amongst which are prevention of HIV infection among women of childbearing age, prevention of unwanted pregnancies among HIV infected women, provision of a comprehensive package of prevention of MTCT of HIV (PMTCT) services, and treatment, care and support to women, children and their families[4]. HIV testing for pregnant women and their partners as well as condom use, and family planning have been the main strategies that have been implemented to achieve the goals of the first two approaches. Knowledge of one’s HIV status through testing accompanied with counseling can lead to the adoption of less risky behavior while anti-retroviral therapy (ART) leads to viral suppression, reducing both heterosexual transmission and MTCT. Indeed, HIV testing amongst pregnant women increased from 54% to 75% between 2012 and 2017 while access to ART also increased from 21.4% to 75.7% during the same period[5–7].

Despite the improvements highlighted in the indicators reported above, the prevalence of HIV amongst pregnant women has remained consistently higher compared with that of the general population and national HIV MTCT rates in HIV-exposed infants has not dropped below the 2% target at 6 weeks since implementation of the eMTCT plan in 2012.

In order to address this disproportionate burden of HIV infection in pregnant women which is serving as barrier to achieving virtual elimination of new pediatric infections, it is imperative to understand the clinical, social and geographical factors associated with this high HIV burden. This will enable the HIV program to focus evidence-based prevention interventions on high-risk women living in the most affected geographical areas, in order to tip the balance in favor of a reduction in HIV infection. An understanding of these risk factors and the implementation of such interventions will also help to bring the country closer to the goal of eliminating pediatric HIV.

## Methods

### Study design and setting

A cross sectional analytic study was conducted in 20 sentinels survey sites across the 10 regions of Cameroon in 2017, which included 60 HIV surveillance health facilities (routine collection points). These health facilities were chosen based on their capacity to provide both ANC and PMTCT services, their location (urban and rural settings in each region of the country), client volume at ANC (capacity to enroll at least 300 pregnant ANC1 attendees during the study period of three months). Urban and rural setting in each region had 3 sites each. At each site, pregnant women aged 15-49 years attending their first ANC were consecutively enrolled until the sample size was reached. The sample size for each region was calculated based on the prevalence of HIV in the region and a desired precision of 95%.

### Study procedure and Participant

Socio-demographic and clinical characteristics were collected from consenting pregnant ANC1 attendees by a nurse without altering the normal functional routine of the health facility. After completing the questionnaire, participants were sent to the laboratory where plasma was collected for HIV and syphilis screening tests according to the routine procedure at the site. After performing HIV and syphilis testing on site, residual plasma was stored in a cryotube at 0 to 8°C, labeled with the subject’s identification code. Twice a month during the study period, a regional supervisor collected samples and questionnaires de-linked them from the corresponding woman and transferred them from the health facility to the central level.

The questionnaires were deposited at the research unit of the National AIDS Control Committee (NACC) and the plasma samples were deposited at the national public health laboratory (NPHL) for analyses.

At the NPHL, HIV screening was performed according to the national serial algorithm for HIV rapid testing (Fig 1). The first test was Determine HIV1/2 (Abbott, Minato-ku Tokyo, Japan). In case this first screening test was reactive, Oraquick (OraSure Technologies, Inc, Bethlehem, Pennsylvania) was used as the confirmatory test. Specimens with indeterminate HIV results at the NPHL were subjected to a tiebreaker test using ImmunoComb^®^ II HIV 1&2 BiSpot. The HIV test results were sent weekly to the NACC to be linked back to the information of the questionnaire through the unique identifier to create a full data set.

**Fig 1.**
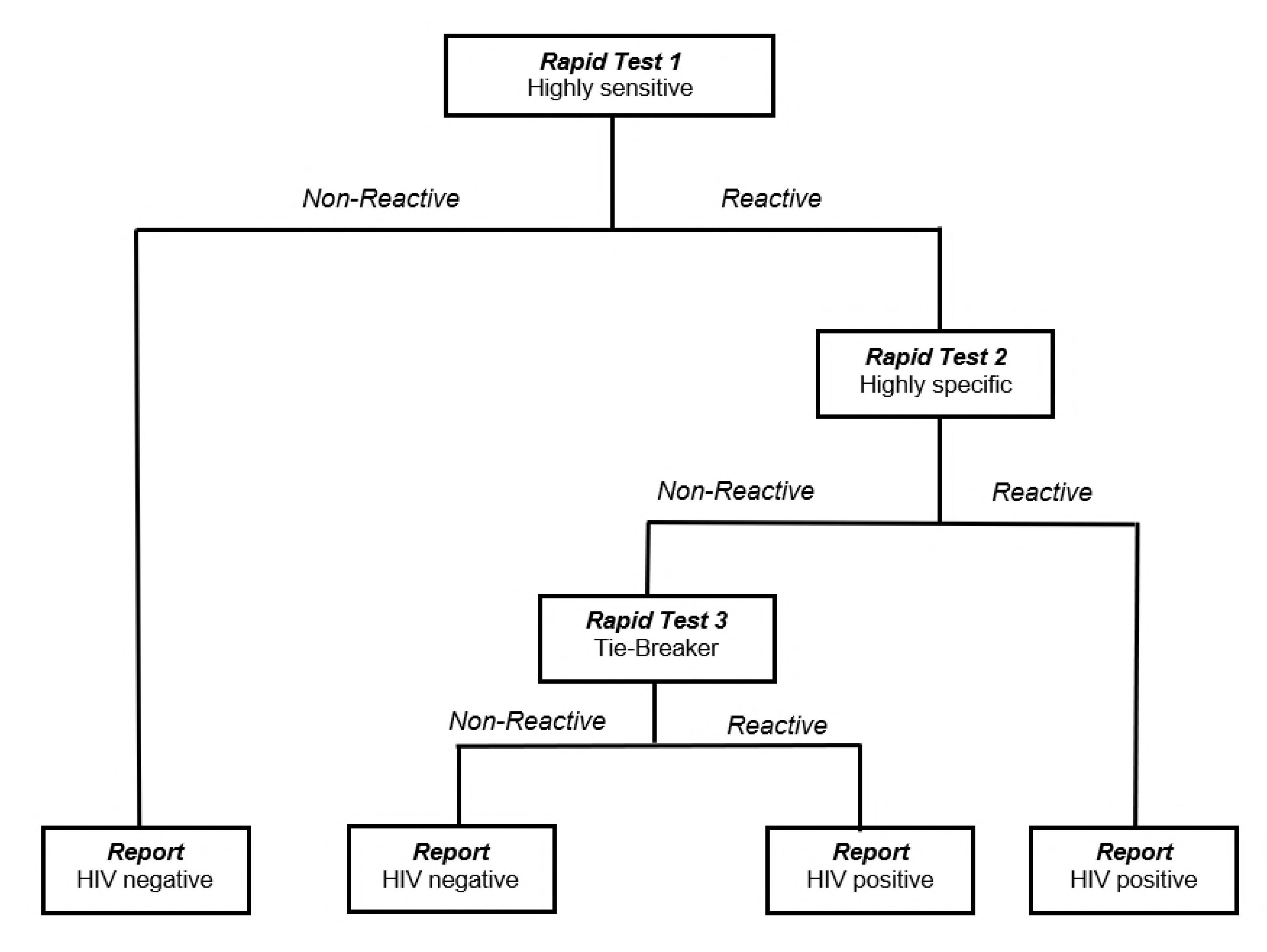
National Algorithm for HIV Rapid Testing

### Statistical analyses

Data was entered in a computer using the CSPRO statistical software. The analysis was performed using STATA/SE version 13.0 (STATACORPS, Texas, USA). Continuous variables were reported as medians with 25th and 75th percentiles, and as means and standard deviations, while categorical variables were described as frequencies and percentages. A multivariate logistic regression was used to investigate the factors associated with HIV infection in pregnant ANC1 attendees. All variables that were significant at 5% in bivariate analysis were introduced into the multivariate model. P values < 0.05 was considered statistically significant.

### Ethical consideration

Ethical clearance for the study was obtained from the Cameroon National Ethics Committee for research. Participation was voluntary without any incentive. HIV tests were offered for free and all women tested positive were placed on ART according to national guidelines. Confidentiality and privacy of the study subjects was ensured by permanently delinking personal identifiers with subject information.

## Results

### Sociodemographic characteristics

A total of 6 859 pregnant women were enrolled in the study. The number of participants varied from 619 in the littoral region to 712 in the North and South-West regions. The mean age was 26.2±6.2 years and young women less than 25 represented 42.7% (2 929/6 859) of the sample with up to 15.1% (1033/6 859) below 20 years of age. Over four fifths (86.2%) attended at least the primary school and 13.5% (949/6859) had university level of education. About 57.3% participants were enrolled in urban area, and almost half (49.4%) were housewife (unemployed) and 17.0% were students (Table 1).

**Table 1:**
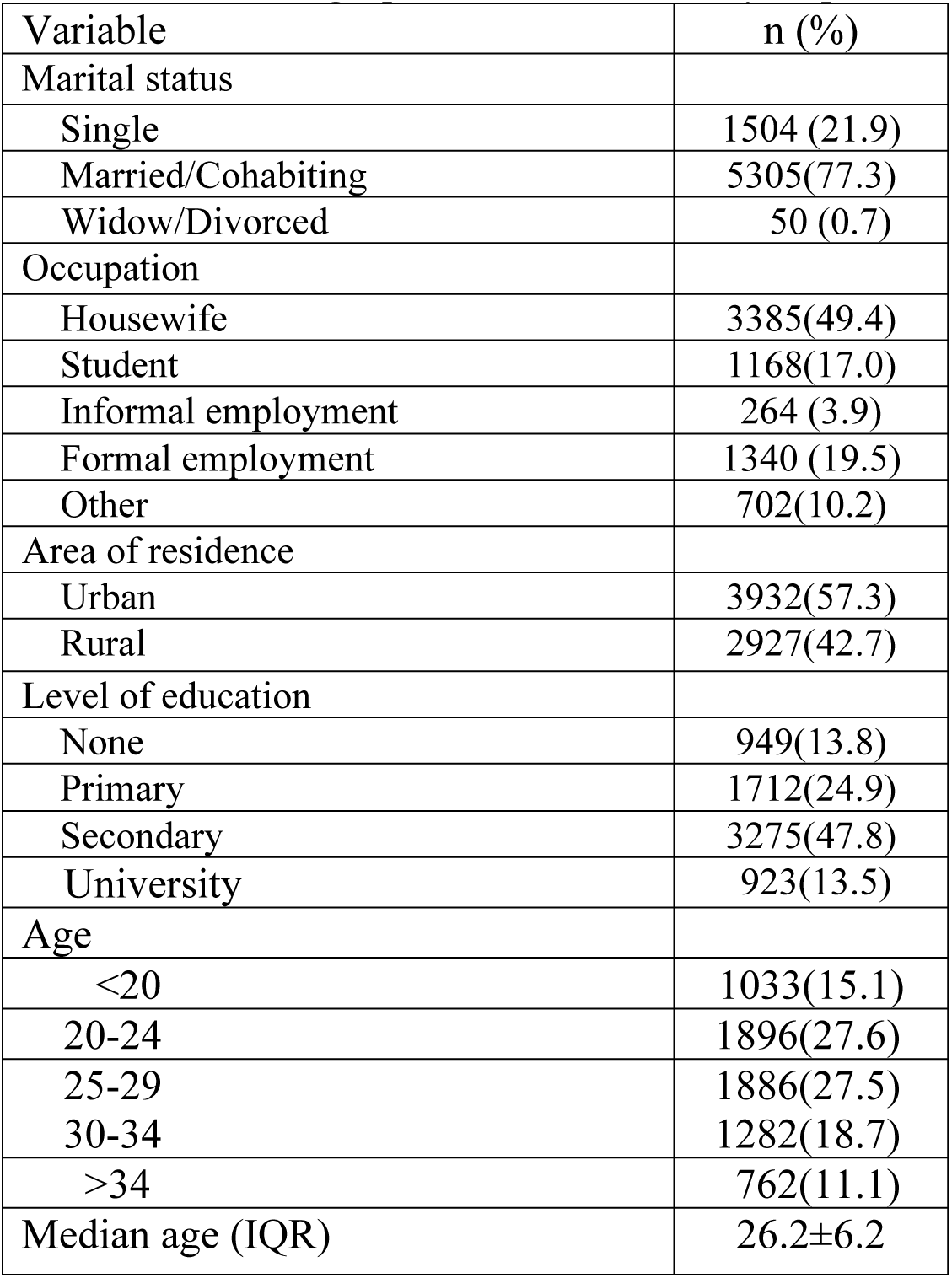
Sociodemographic characteristics of respondents

| Variable | n (%) |
| --- | --- |
| Marital status |  |
| Single | 1504 (21.9) |
| Married/Cohabiting | 5305(77.3) |
| Widow/Divorced | 50 (0.7) |
| Occupation |  |
| Housewife | 3385(49.4) |
| Student | 1168(17.0) |
| Informal employment | 264 (3.9) |
| Formal employment | 1340 (19.5) |
| Other | 702(10.2) |
| Area of residence |  |
| Urban | 3932(57.3) |
| Rural | 2927(42.7) |
| Level of education |  |
| None | 949(13.8) |
| Primary | 1712(24.9) |
| Secondary | 3275(47.8) |
| University | 923(13.5) |
| Age |  |
| <20 | 1033(15.1) |
| 20-24 | 1896(27.6) |
| 25-29 | 1886(27.5) |
| 30-34 | 1282(18.7) |
| >34 | 762(11.1) |
| Median age (IQR) | 26.2±6.2 |

### HIV prevalence and associated risk factors

The overall prevalence of HIV was 5.7% (95% CI: 5.1 – 6.2). The highest was recorded in the Centre (9.6%), East (9.7%) and South-West (9.0%) regions, while the lowest was recorded in the North region [2.6% (19/700)]. The HIV infection is significantly corelated with the employment status. The odds of HIV were almost two time among employee of informal sector compared to housewife [OR=1.83 (95% CI:1.36 – 2.51)]. Education level was also significantly related to HIV infection, the women who reported having achieved primary school education [OR=2.26 (95%CI:1.49-3.28)] and secondary school education [OR=1.71(95%CI:1.16-2.47)] had higher prevalence as compared to those who reported never having attended school. The odds of being infected with HIV was higher among women residing in the Center [OR=3.15(95%CI:1.19-5.17)], East [OR=3.20(95%CI:1.97-5.24)], Nord-West [OR=2.02(95%CI:1.19-3.41)] and South-West [OR=2.96((95%CI:1.80-4.85)] regions of the country. The HIV infection increase with age group. The odds of HIV infection was more than three times among women age above 20 years as compared with those aged less than 20 years (20-24 years [OR=3.03(95%CI:1.76-5.18)], 25-29 years [OR=3.86(95%CI:2.27-6.57)], 30-34 years [OR=5.61(95%CI:3.29-9.56)], age >34 years [OR=7.13(95%CI:4.13-12.28)]). Multiparous are almost three time more at risk of infection than nulliparous [OR=2.79 (95%CI: 2.01-3.88)] and the odds of infection are even higher among grand multiparity (4 and more deliveries) [OR=1.47 (95%CI:1.81-1.82).

Being married or cohabiting ([OR=0.76 (95% CI= 0.59 – 0.96)]) as well as being a student ([OR=0.45 (95%CI:0.30 – 0.69)]) were found to be associated with lower odds HIV infection. There was no statistical association between HIV infection and the area of residence (residing in rural or urban areas) (Table 2).

**Table 2:** Factors associated with HIV infection amongst pregnant women

| Factors | #HIV- (%) | #HIV+ (%) | OR (95% CI) | P-value |
| --- | --- | --- | --- | --- |
| Marital status |  |  |  |  |
| Single | 1403(93.3) | 101(6.7) | 1 |  |
| Married/Cohabiting | 5019(94.6) | 286(5.4) | 0.76(0.59-0.96) | 0.022 |
| Widowed/Divorced | 44(88.0) | 6(12.0) | 2.54(0.73-8.86) | 0.144 |
| Occupation |  |  |  |  |
| Housewife | 3217(95.0) | 168(5.0) | 1 |  |
| Student | 1141(97.7) | 27(2.3) | 0.45(0.30-0.69) | 0.000 |
| Formal employee | 245(92.8) | 19(7.2) | 1.43(0.92-2.47) | 0.115 |
| Informal employee | 1223(91.3) | 117(8.7) | 1.83(1.36-2.51) | 0.000 |
| Others | 640(91.2) | 62(8.8) | 1.85(1.37- 2.51) | 0.000 |
| Area of residence |  |  |  |  |
| Urban | 3711(94.4) | 221(5.6) | 1 |  |
| Rural | 2755(94.1) | 172(5.9) | 1.05 (0.86-1.29) | 0.652 |
| Level of education |  |  |  |  |
| None | 916(96.5) | 33(3.5) | 1 |  |
| Primary | 1582(92.5) | 129(7.5) | 2.26(1.49-3.28) | 0.000 |
| Secondary | 3084(94.2) | 191(5.8) | 1.71(1.16-2.47) | 0.005 |
| University | 883(95.7) | 40(4.3) | 1.26(0.78-2.01) | 0.340 |
| Age |  |  |  |  |
| <20 | 1017(98.4) | 16(1.6) | 1 |  |
| 20-24 | 1810(95.5) | 86(4.5) | 3.02(1.76-5.18) | 0.000 |
| 25-29 | 1730(94.3) | 105(5.7) | 3.86(2.27-6.57) | 0.000 |
| 30-34 | 1178(91.9) | 104(8.1) | 5.61(3.29-9.56) | 0.000 |
| >34 | 731(89.9) | 82(10.1) | 7.13(4.13-12.28) | 0.000 |
| Nulliparous |  |  |  |  |
| Yes | 1587(97.5) | 41(2.5) | 1 |  |
| No | 4879(97.3) | 352(6.7) | 2.79(2.01-3.88) | 0.000 |
| Multiparous |  |  |  |  |
| 1 -3 deliveries | 2489(94.5) | 146(5.5) | 1 |  |
| 4 and more deliveries | 2390(92.1) | 206(7.9) | 1.47(1.18-1.82) | 0.001 |
| Region |  |  |  |  |
| Adamawa | 655(96.7) | 22(3.3) | 1 |  |
| Centre | 614(90.4) | 65(9.6) | 3.15(1.91-5.17) | 0.000 |
| East | 632(90.3) | 68(9.7) | 3.20(1.97-5.24) | 0.000 |
| Far-North | 658(96.3) | 25(3.7) | 1.13(0.63-2.02) | 0.671 |
| Littoral | 597(96.4) | 22(3.6) | 1.09(0.60-2.00) | 0.758 |
| North | 700(97.4) | 19(2.6) | 0.81(1.19-3.41) | 0.503 |
| North-West | 634(93.6) | 43(6.4) | 2.02(1.19-3.41) | 0.009 |
| South | 639(94.7) | 36(5.3) | 1.68(0.97-2.88) | 0.098 |
| South-West | 654(91.0) | 65(9.0) | 2.96(1.80-4.85) | 0.000 |
| West | 683(96.1) | 28(3.9) | 1.22(0.69-2.15) | 0.489 |

### Multivariate analysis

Multivariate logistic regression analysis revealed that age is significantly associated with HIV infection. (Women aged 20-24 years [aOR=2.8 (95% CI:1.6-4.9)] and those aged 25–29 years [aOR=3.1 (95% CI:1.7-5.7)] are three times more likely to be infected than those less than 20 years of age Those aged 30-34 years and > 34 years are four [aOR=2.9(95%CI:1.5-5.6)] and five [aOR=4.2(95%CI:2.2-8.72)] times more likely to be infected as compared to those less than 20). Multiparity is also associated to the risk of infection [aOR=1.5(95%CI:1.1-2.2)] as well as living in the Centre [aOR=2.6(95%CI:1.5-4.4)], East [aOR=3.2(95%CI:1.9-5.2)] and South-West [aOR=2.4(95%CI:1.4-4.0)] regions compared to living to Adamawa. However, married [aOR=0.5(95%CI:0.4-0.7)] compared single and student [aOR=0.5(95%CI:0.3-0.8)] compared to housewife were inversely associated with HIV. infection. No significant interaction with level of education was observed (Table 3).

**Table 3:** Predictors of HIV infection amongst pregnant women

| Variables | aOR (95% CI) | P-value |
| --- | --- | --- |
| Marital status |  |  |
| Single | 1 |  |
| Married/Cohabiting | 0.5(0.4-0.7) | 0.000 |
| Widowed/Divorced | 0.8(0.3-2.0) | 0.621 |
| Occupation |  |  |
| Housewife | 1 |  |
| Student | 0.6(0.3-0.9) | 0.021 |
| Informal employee | 1.2(0.7-2.0) | 0.577 |
| Formal employee | 1.3(0.9-1.6) | 0.099 |
| Others | 1.3(0.9-1.0) | 0.096 |
| Nulliparous |  |  |
| Yes | 1 |  |
| No | 1.5(1.1-2.2) | 0.029 |
| Age |  |  |
| <20 | 1 |  |
| 20-24 | 2.8(1.6-4.9) | 0.000 |
| 25-29 | 3.1(1.7-5.7) | 0.000 |
| 30-34 | 4.3(2.3-7.9) | 0.000 |
| >34 | 5.7(3.1-10.8) | 0.000 |
| Level of instruction |  |  |
| None | 1 |  |
| Primary | 1.4 (1.0-2.3) | 0.072 |
| Secondary | 1.2(0.8-1.9) | 0.406 |
| University | 0.8(0.4-1.3) | 0.345 |
| Region |  |  |
| Adamawa | 1 |  |
| Centre | 2.6(1.5-4.4) | 0.000 |
| East | 3.2(1.9-5.2) | 0.000 |
| Far-North | 1.2(0.7-2.2) | 0.561 |
| Littoral | 0.9(0.5-1.6) | 0.646 |
| North | 0.9(0.5-1.6) | 0.668 |
| North-West | 1.7(1.0-3.0) | 0.050 |
| South | 1.5(0.9-2.6) | 0.105 |
| South-West | 2.4(1.4-4.0) | 0.001 |
| West | 1.1(0.7-1.9) | 0.838 |

## Discussions

The overall goal of this study was to assess the factors associated with HIV infection among pregnant women in Cameroon. The national HIV prevalence was estimated in this vulnerable population and then risk factors associated with infection, were analyzed. The results of the study show that HIV infection among pregnant women is decreasing. In 2012, the prevalence of HIV in this group was 7.8% (n=6521, 95% CI = [7.15% - 8.45%]) while our results show a prevalence of 5.7% (n=6859, 95% CI = [5.17% - 6.27%]) [3, 8]. This decrease is the result of several strategies implemented by the country to strengthen the prevention of HIV particularly amongst women who are disproportionately affected by the virus. Some of these strategies include implementation of the four pillars of the eMTCT plan [4], PMTCT integration into ANC, rapid scale up of PMTCT coverage nationwide as well as implementation of Option B+. Indeed, condom use increased between 2012 to 2017, the number of clinics offering PMTCT services increased by 63% (from 1600 to 4342) between 2012 and 2017. Since 2014, option B+ was adopted by the country as the national PMTCT strategy. The HIV infected women are given a single pill of ART combination (Tenofovir+Lamivudine+Efavirenz) per day and are kept on treatment after the delivery. The efficacity of the prevention program can be confirmed by the high HIV testing acceptance rate of almost 99% at ANC. In addition, the reduction of HIV prevalence can be further confirmed by the low prevalence registered among adolescent women in this study. In 2012, the HIV prevalence among pregnant women under 20 years of age was 3.4% while in our study, it was 1.6% [8].

Our results showed that single women are two time more likely to be infected by HIV than those who are married or cohabiting. Others study had reported similar findings that peoples engaged in a marital relationship are less exposed to HIV infection[9] This can be explained by the fact that individuals are more likely to have multiple partners before marriage or after divorce or separation[10, 11]. Moreover, due to their economic vulnerability, single women are more involved in transactional sex which increases their exposure to HIV as a result [12]. The higher odds of infection in divorced and widowed women compared to those that are married could be due to the fact that the positive HIV status of one partner might contribute to separation and divorce while widowhood include women who have lost their partners due to HIV [10, 13]

The odds of being infected with HIV increased with age. Women aged 25 years and above are at least two times more at risk of being infected with HIV compared with those that are younger. This result is in accordance with the distribution of HIV infection in the general population, were the prevalence of HIV is higher in the higher age-groups[3]. The age at first intercourse is below 15 years for many women in Cameroon [14]. This results in a progressively increasing duration of exposure to sexual activity and thus more risk of infection by HIV.

In addition, the odds of HIV infection increased with gravidity. Higher gravidity implies more exposure to unprotected sex, hence higher risk of being infected with the virus. As we stated before, the improvement registered in the implementation of prevention programs particularly among women has certainly contributed to reduce new infection, with primigravid women newly enrolled into ANC care having less odds of being infected with HIV than multi-gravid women. Beyond the desire of pregnancy, higher numbers of previous pregnancies can also reflect inadequate power of women to negotiate protected sex. This has been demonstrated in many studies to be a risk factor of HIV infection[12]. This result provides evidence for the need of interventions to control HIV, focusing on family planning and the promotion of condom use during sex for women, especially those that are widowed and divorced.

Pregnant women living in the Centre, East, North-West and South-West regions were more at risk of HIV infection than those in the other regions. These four regions are the most affected in the country after the South region which has the highest HIV prevalence [3]. These regions should thus be considered for priority interventions to curb this high HIV burden. Compared with the prevalence obtained in previous studies conducted in 2012, the prevalence in the South region has dropped from 7.5% to 5.1% and in the Littoral, the drop was even more important, (from 8.8% to 3.6%).

The bivariate analysis showed an association between level of education and HIV infection amongst pregnant women, as many other studies have [11, 15]. But the level of education was not significant in multivariate analysis. This finding reassures us that the sensitization carried out on HIV prevention, care and treatment has been efficient. Indeed, between 2011 and 2014 number of women in Cameroon who have proper knowledge about HIV as reported in the demographic health survey (2011) and in the Multi indicator cluster survey (2014) has increased from 26% to 30.1%.[2, 14].

The strength of our study is the large population size and the fact that it covers all ten regions of the country offers the possibility to generalize the results. A possible limitation is the limited number of factors considered in the study. The economic characteristics as well as the cultural environment of participants, which have been shown in previous studies to be significantly related to HIV infection, were not collected in this study. Nevertheless, the general socio-cultural and economic situation of the different regions which have been well established in previous studies were considered in explaining the results of the study.

## Conclusion

Our results showed that even though HIV infection among pregnant women is decreasing, this group is still disproportionately affected as the HIV prevalence is still higher than that of the general population. For the HIV prevention program amongst pregnant women to be effective, it must specifically target older pregnant women, who are single or divorced and having a higher gravidity. Special attention should be paid Centre, South-West, North-West and East regions where comprehensive prevention strategies should be undertaken.

## Acknowledgements

We are grateful to pregnant women who provided consent to participate in this survey. We also thank the personnel of the PMTCT services and of the laboratories of all the participating health facilities. We also thank the Ministry of Public Health for providing support for the implementation of the study throughout the country.

## Contributors

Study design: JDA, JF, ELG, SCB, CKN, DGT, AM and JBEN.

Investigation: JDA, ELG, CKN, JF, DGT, AM and YM

Coordination: JBEN, LB, SCB and JDA

Funding: JBEN

Data analysis: JDA and CKN

Writing original draft: JDA and ELG

All authors revised the report critically and approved the final version.

## Conflict of Interest

JF received consultancy honorarium for the sentinel survey in 2016.The others have no other conflict of interest to declare.

## References

[1] UNAIDS. http://www.unaids.org/sites/default/files/media_asset/UNAIDS_FactSheet_en.pdf, july 2018

[2] Institut National de la Statistique (INS) et ICF. International. “ Enquête Démographique et de Santé et à Indicateurs Multiples du Cameroun 2011”. Calverton, Maryland, USA: INS et ICF International. 2012. https://dhsprogram.com/pubs/pdf/fr260/fr260.pdf

[3] Comité National de Lutte contre le Sida. “Surveillance sentinelle du VIH et de la Syphilis chez les femmes enceintes en consultation prénatale au Cameroun en 2016.”.2017.

[4] Ministère de la santé Publique, “Plan d’Elimination National de la Transmission Mère Enfant du VIH à l’horizon 2015.” 2012.

[5] Comité National de Lutte contre le Sida. “Rapport progrès PTME N° 12” 2017.

[6] Comité National de Lutte contre le Sida. “Rapport Annuel 2012 des Activité de lutte contre le VIH/Sida et les IST au Cameroun.” http://cnls.cm/sites/default/files/rapport_annuel_cnls_2012.pdf.

[7] Comité National de Lutte contre le Sida. “Rapport Annuel 2016 des Activité de Lutte Contre le VIH, le Sida et les IST au Cameroun.”. http://cnls.cm/sites/default/files/rapport_annuel_cnls_2016.pdf.

[8] Comité National de lutte contre le Sida. “Etude de faisabilité d’un système de surveillance sentinelle du VIH basé sur les données PTME du Cameroun au Cameroun”. Avril 2013.

[9] M. Mabaso, Z. Sokhela, N. Mohlabane, B. Chibi, K. Zuma, and L. Simbayi, “Determinants of HIV infection among adolescent girls and young women aged 15–24 years in South Africa: a 2012 population-based national household survey,” BMC Public Health, vol. 18, no. 1, Dec. 2018.

[10] L. A. Chimoyi and E. Musenge, “Spatial analysis of factors associated with HIV infection among young people in Uganda, 2011,” BMC Public Health, vol. 14, no. 1, Dec. 2014.

[11] B. A. Mohamed and M. S. Mahfouz, “Factors Associated with HIV/AIDS in Sudan,” BioMed Research International, 2013. [Online]. Available: https://www.hindawi.com/journals/bmri/2013/971203/. [Accessed: 18-Aug-2018].

[12] R. C. Dellar, S. Dlamini, and Q. A. Karim, “Adolescent girls and young women: key populations for HIV epidemic control,” J Int AIDS Soc, vol. 18, no. 2 Suppl 1, Feb. 2015.

[13] E. V. Boisson, “Factors associated with HIV infection are not the same for all women,” Journal of Epidemiology & Community Health, vol. 56, no. 2, pp. 103–108, Feb. 2002.

[14] Institut National de la Statistique. 2015. “Enquête par grappes à indicateurs multiples (MICS5)”, 2014, Rapport Final. Yaoundé, Cameroun, Institut National de la Statistique.

[15] Swai et al. “Surveillance of HIV and syphilis infections among antenatal clinic attendees in Tanzania-2003/2004.” BMC Public Health, vol. 6, no 1, pp. 91, Apr. 2016, https://bmcpublichealth.biomedcentral.com/articles/10.1186/1471-2458-6-91.

